# Effect of SSRI discontinuation on anxiety-like behaviours in mice

**DOI:** 10.1101/2021.05.29.446266

**Authors:** Helen M Collins, Raquel Pinacho, Dersu Ozdemir, David M Bannerman, Trevor Sharp

## Abstract

**Background:** Abrupt cessation of therapy with a selective serotonin reuptake inhibitor (SSRI) is associated with a discontinuation syndrome, typified by numerous disabling symptoms including anxiety. Surprisingly little is known of the behavioural effect of SSRI discontinuation in animals.

**Aim:** Here, the effect of SSRI discontinuation on anxiety-like behaviour was systematically investigated in mice.

**Methods:** Experiments were based on a 3-arm experimental design comprising saline, continued SSRI and discontinued SSRI. Mice were assessed 2 days after SSRI discontinuation over a 5 day period using the elevated plus maze (EPM) and other anxiety tests.

**Results:** An exploratory experiment found cessation of paroxetine (12 days) was associated with decreased open arm exploration and reduced total distance travelled, in male but not female mice. Follow-up studies confirmed a discontinuation effect on the EPM in male mice after paroxetine (12 days) and also citalopram (12 days). Mice receiving continued paroxetine (but not citalopram) also showed decreased open arm exploration but this was dissociable from effects of discontinuation. The discontinuation response to paroxetine did not strengthen after 28 days treatment but was absent after 7 days treatment. A discontinuation response was not decernable in other anxiety and fear-learning tests applied 3-5 days after treatment cessation. Finally, discontinuation effects on the EPM were typically associated with decreased locomotion on the test. However, separate locomotor testing implicated anxiety-provoked behavioural inhibition rather than a general reduction in motor activity.

**Conclusion:** Overall, the current study provides evidence for a short-lasting behavioural discontinuation response to cessation of SSRI treatment in mice.

## Introduction

Selective serotonin (5-hydroxytryptamine; 5-HT) reuptake inhibitors (SSRIs) are currently the first-line pharmacological treatment for major depression and anxiety disorder. Despite their generally improved tolerability and side-effect profile compared to older antidepressants (Cipriani et al., 2018), the abrupt cessation of SSRI treatment, like other antidepressants, is often associated with a disabling discontinuation syndrome (Haddad, 1997). Typical SSRI discontinuation symptoms include heightened anxiety, insomnia, nausea, dizziness, irritability and increases in suicidal thoughts (Delgado, 2006; Warner et al., 2006; Horowitz and Taylor, 2019). SSRI discontinuation is considered distinct from depression relapse in that symptoms appear within days of discontinuation, whereas relapse typically takes several weeks to manifest. Moreover, discontinuation symptoms are often different to those evident during the depressive episode and include the emergence of somatic symptoms (Delgado, 2006). Problematic effects of SSRI discontinuation are reported by depressed patients as well as other patient groups including those with seasonal affective disorder, social anxiety and panic disorder (Lader et al., 2004; Montgomery et al., 2005; Black et al., 1993).

SSRI discontinuation has recently risen to prominence following reports that the syndrome may be more common, disabling and longer-lasting that previously recognised (APPG, 2018; Davies and Read, 2019). Despite its clinical importance there are few systematic investigations of SSRI discontinuation and its mechanism remains unknown. As an example of the problem we are aware of just two preclinical studies that directly address the behavioural effects of SSRI discontinuation. One study reported enhanced acoustic startle response in rats 2 days after discontinuation from repeated treatment with citalopram (Bosker et al., 2010). Another study reported increased locomotor behaviour 4 h after the first “missed dose” of repeated fluoxetine in rats, and this effect dissipated within 4 days (Bjork et al., 1998). In addition, although not reported as investigation of SSRI discontinuation itself, a small number of studies report no change in anxiety or locomotor behaviour in rats 1-3 weeks after the last dose of repeated SSRI administration (Elizalde et al., 2008; Popa et al., 2010; Bouet et al., 2012; Strekalova et al., 2013). Taken together, these limited preclinical findings suggest short-lasting behavioural changes associated with SSRI discontinuation in rodents but this requires systematic investigation.

It is noteworthy that previous preclinical studies have identified behavioural correlates of withdrawal from a wide variety of drugs including opiates, alcohol and psychostimulants (Emmett-Oglesby et al., 1990; Vuong et al., 2010; El Hage et al., 2012; Perez and De Biasi, 2015). Much of this work has centred on behavioural readouts of anxiety, a core feature of many drug withdrawal states (note that the term ‘withdrawal’ as opposed to ‘discontinuation’ is commonly used for drugs associated with compulsive drug seeking behaviour, which is not the case for SSRIs or other antidepressants). Such work led to novel insights into drug withdrawal mechanisms and aided the identification of treatment strategies, as exemplified by the use of clonidine to manage anxiety and other symptoms of opiate withdrawal (Gowing et al., 2002; Kosten and Baxter, 2019).

The present study systematically investigated the effect of SSRI discontinuation on the behaviour of mice, with a focus on anxiety which is a core symptom of SSRI discontinuation in patients. Also, both anxiety and fear are well known to be modulated by pharmacological and genetic manipulations of the 5-HT transporter (Handley and McBlane, 1993; Line et al., 2011; Barkus et al., 2014; Lima et al., 2019). Paroxetine was selected for detailed study since discontinuation from this SSRI is considered particularly problematic in patients (Fava et al., 2015), likely due in part to its short half-life (Price et al., 1996). Paroxetine treatment duration and frequency were varied to optimise conditions to detect discontinuation effects. Citalopram was also added for comparison due to its greater selectivity for the 5-HT transporter than paroxetine and its short half-life in rodents (Fredricson Overø, 1982). Anxiety-like behaviours were measured using a battery of tests including the elevated plus maze (EPM), which has proven sensitive in detecting the withdrawal states of other drugs (Emmett-Oglesby et al., 1990; Vuong et al., 2010; El Hage et al., 2012; Perez and De Biasi, 2015), and is highly sensitive to manipulations of the 5-HT system (Briley et al., 1990; Handley and McBlane, 1993; Ohmura et al., 2020).

## Materials & Methods

### Animals

Adult mice (C57BL/6J, 8-10 weeks, Charles River) were housed (21°C on a 12 h light/dark cycle; lights off 19:00 to 7:00) with littermates in open-top cages (3-6 mice per cage) lined with sawdust bedding, for at least 1 week before the start of treatment. Food and water were available *ad libitum*. Cardboard sizzle nests were used for cage enrichment, and mice were handled using a cardboard tunnel to minimize stress associated with repeated injections (Gouveia and Hurst, 2019). Experiments followed the principles of the ARRIVE guidelines and were conducted according to the U.K. Animals (Scientific Procedures) Act of 1986 with appropriate personal and project licence coverage.

### Drugs and reagents

Paroxetine hydrochloride (Abcam, ab120069; Apexbio, B2252-APE) and citalopram hydrobromide (Abcam, ab120133) were dissolved in saline (1 and 2 mg/ml, respectively). Drug dose (10 mg/kg s.c.) was chosen on the basis of both the half-life of paroxetine (6.3 h) and citalopram (1.5 h) in rodents (Kreilgaard et al., 2008) (Fredricson Overø, 1982), and previous studies reporting antidepressant actions of repeated administration of these drugs in mice (Elizalde et al., 2008). Treatment frequency (once- or twice-daily) and duration (7-28 days) were varied in an attempt to optimise the detection of discontinuation effects.

### Experimental design

Mice were allocated to 1 of 3 experimental groups by stratified randomisation: i) saline group: saline + saline continuation; ii) continuation group: SSRI + SSRI continuation; iii) discontinuation group: SSRI + saline continuation. Mice were weighed daily beginning 2 days before the start of the experiment to establish a baseline weight, and habituate mice to handling. Mice received once- or twice-daily injections of paroxetine, or the equivalent volume of saline, for 7, 12 or 28 days, then treatment was either continued (saline and continuation groups) or swapped to saline injections (discontinuation group) for a further 5 days during which behavioural tests were conducted. The effect of twice-daily injections of citalopram or saline for 12 days was also tested using a similar experimental design. Testing commenced either 18 h (once-daily) or 6 h (twice-daily) after the last injection. The 5 day discontinuation period was based on previous rodent experiments reporting discontinuation effects (Bosker et al., 2010; Trouvin et al., 1993) and aimed to capture the timeframe when paroxetine and citalopram fall to undetectable levels in the blood plasma (Benmansour et al., 1999; Cremers et al., 2000). An initial exploratory investigation of paroxetine used mixed sex groups (6 males, 6 females per group) and revealed sexually dymorphic effects. Consequently, all follow up experiments utilised male mice. Details of this and all other experiments are summarised in *Table 1*.

**Table 1.**
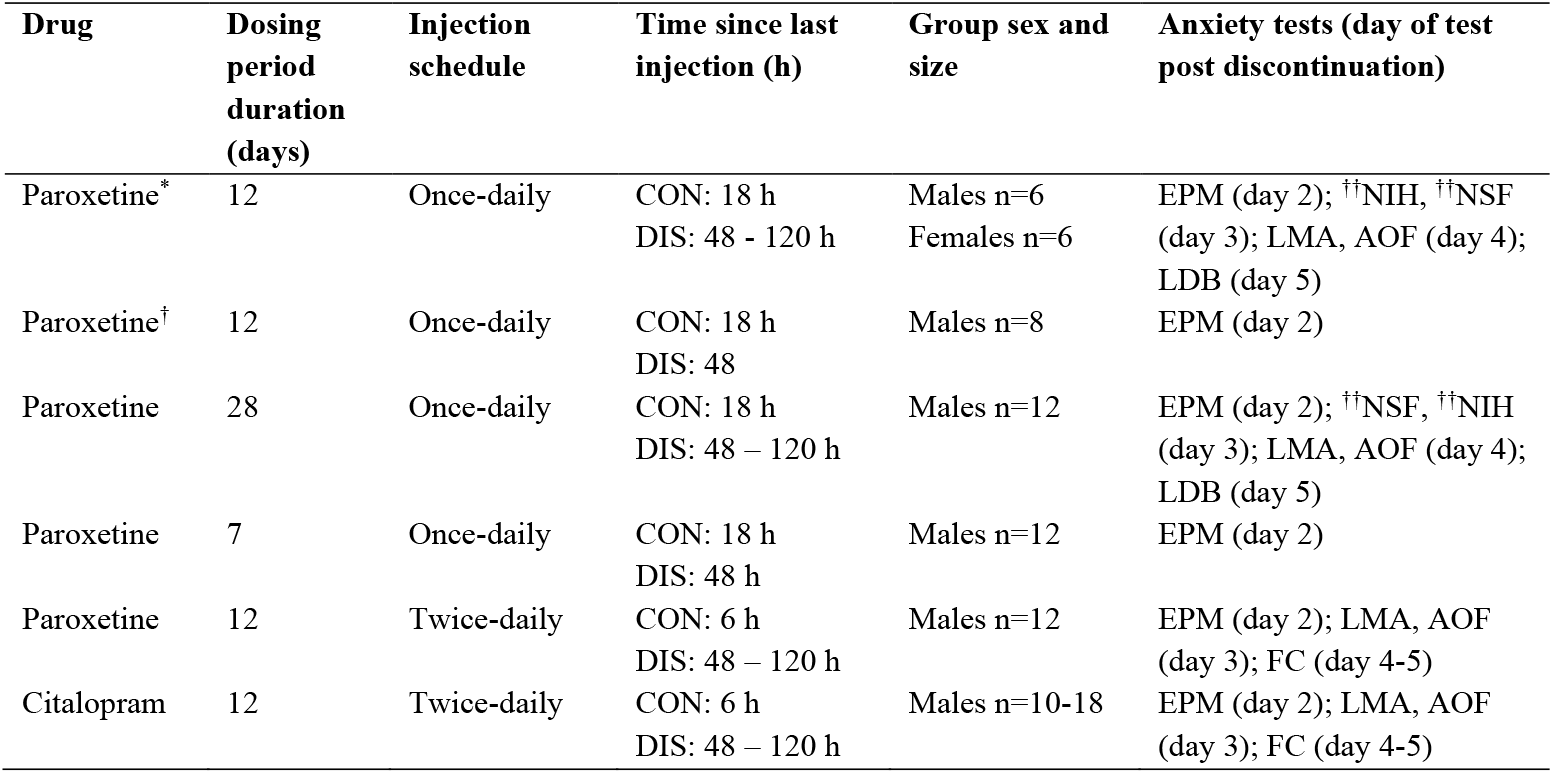
Experimental design. Elevated plus maze, EPM; locomotor activity, LMA; aversive open field, AOF; light/dark box, LDB; fear conditioning, FC. Time since last injection of SSRI for Continuation (CON) and Discontinuation (DIS) groups, DIS receive saline injections on discontinuation days. *Exploratory experiment, †data were pooled with the exploratory experiment, ††data excluded from the study (see Methods for details).

| Drug | Dosing period duration (days) | Injection schedule | Time since last injection (h) | Group sex and size | Anxiety tests (day of test post discontinuation) |
| --- | --- | --- | --- | --- | --- |
| Paroxetine* | 12 | Once-daily | CON: 18 h<br>DIS: 48 - 120 h | Males n=6<br>Females n=6 | EPM (day 2); ††NIH, ††NSF (day 3); LMA, AOF (day 4); LDB (day 5) |
| Paroxetine† | 12 | Once-daily | CON: 18 h<br>DIS: 48 | Males n=8 | EPM (day 2) |
| Paroxetine | 28 | Once-daily | CON: 18 h<br>DIS: 48 – 120 h | Males n=12 | EPM (day 2); ††NSF, ††NIH (day 3); LMA, AOF (day 4); LDB (day 5) |
| Paroxetine | 7 | Once-daily | CON: 18 h<br>DIS: 48 h | Males n=12 | EPM (day 2) |
| Paroxetine | 12 | Twice-daily | CON: 6 h<br>DIS: 48 – 120 h | Males n=12 | EPM (day 2); LMA, AOF (day 3); FC (day 4-5) |
| Citalopram | 12 | Twice-daily | CON: 6 h<br>DIS: 48 – 120 h | Males n=10-18 | EPM (day 2); LMA, AOF (day 3); FC (day 4-5) |

### Behavioural tests

Mice were habituated to a holding room (i.e. not their home cage room) for at least 1 h before testing and on the 3 days preceding behavioural testing. All testing was conducted in the light phase (10:00 – 17:00 h) by an observer blind to treatment. Other than the changes in anxiety-related measures reported herein, no overt behavioural or somatic effects to continued/discontinued SSRI treatment were observed. Both a novelty-induced hyponeophagia test (Line et al., 2011) and novelty-supressed feeding test (Santarelli et al., 2003) were conducted on day 3 following discontinuation from 12- and 28-day once-daily paroxetine treatment but results were confounded by an effect of test order, and hence data were excluded from the study.

#### Elevated Plus Maze (EPM)

The EPM model assesses approach-avoidance behaviour relying on the conflict between the innate aversion of open, elevated spaces and the exploratory drive in mice (Pellow et al., 1985; Komada et al., 2008). EPM experiments were carried out as previously described for studies examining the role of the 5-HT transporter in unconditioned anxiety (Line et al., 2011). The EPM (50 cm off the floor) was placed in a dimly-lit room and comprised 2 open arms (35 × 6 cm) perpendicular to 2 closed arms (35 × 6 cm, 20 cm walls), with a central region (6 × 6 cm). Mice were initially placed facing the walls at the far end of the closed arm, and movement was automatically tracked for 300 s (ANY-maze software, Stoelting Co.). Key parameters were time spent in the open arms, open arm entries and latency to enter open arms with distance travelled being used as the principal readout of locomotor activity (closed arm entry was also recorded).

#### Locomotor activity (LMA)

Given the potential confound of changes in locomotion on anxiety measures, previous studies highlight the value of assessing locomotion in separate low anxiety environments (Lister, 1990). Here, spontaneous LMA was assessed in a dimly-lit room as previously described (Line et al., 2011). Mice were placed individually into an unfamiliar plastic cage (42 × 22 × 20 cm) lined with sawdust and covered with a perforated plexiglass lid. LMA was monitored by horizontal and vertical infrared beams, and the number of beam breaks was automatically recorded in 5 min bins for 60 min (Photobeam Frame software, San Diego Instruments).

#### Aversive open field (AOF)

The open field test assesses exploration and behaviour in an anxiogenic environment (Prut & Belzung, 2003). Mice were placed in a brightly-lit, white cylindrical chamber (30 cm radius) and movement was automatically tracked for 600 s (ANY-maze software). The centre zone was designated as a central circle of 10 cm radius. Key parameters were time spent in the centre zone and distance travelled.

#### Light/dark box (LDB)

The LDB assesses approach-avoidance conflict in rodents based on their innate fear of brightly-lit places (Bourin & Hascoet, 2003). The LDB arena (Pritchett et al., 2015) was inside a sound-attenuating cubicle and consisted of a dark, enclosed compartment (21 cm long x 16 cm high x 16 cm wide) separated by a small doorway from a brightly-lit, open compartment (46.5 cm long x 21 cm high x 21 cm wide). Mice were placed in a corner of the dark compartment and their movement was automatically tracked for 600 s by horizontal infrared beams (activity monitor software, Med Associates Inc). Key parameters were time spent in light zone and distance travelled.

#### Fear conditioning

Assessment of fear learning was performed as described previously (Line et al., 2014) using conditioning chambers (Med Associates Med Associates Inc., USA) with a floor of metal bars connected to a scrambled-shock generator (controlled by Med-PC IV software programme). In brief, mice were placed individually in a covered plastic box for 30 min before each session. Two different conditioning contexts were used, each associated with a specific scent and distinctive wall layout (black and white striped walls, lavender essential oil *versus* grey walls, sandalwood essential oil). Mice were trained in one context and tested in the other (context was counterbalanced across training and test days). On the training day, after a 180 s acclimatisation period, mice received 2 tone-shock pairings, each consisting of a 30 s tone (72 dB, 2900 Hz) paired with a shock (0.5 ms, 0.3 mA) delivered in the final 0.5 s of the tone (180 s inter-trial interval). On the test day, 24 h later, mice received the same tone presentations but did not receive the shock. Freezing was determined from video recordings (analysed by NIH ImageJ with a customised script) and defined as <0.07% pixel change in two consecutive frames (1 frame per second). Pre-tone freezing was calculated as % freezing in the 30 s before the tone, and freezing to tone was calculated as the % freezing for 30 s of tone. The change in freezing levels (ΔFreezing) was then calculated [freezing to tone – pre-tone freezing] and presented as average ΔFreezing per day.

### Statistical analysis

D’Agostino-Pearson’s test for normality was applied to all data sets. If data were normally distributed, then one-way ANOVA was used with Fisher’s Least Significant Difference (LSD) to compare treatments. If data were not normally distributed, Kruskal-Wallis with Fisher’s LSD was used. Sex was assessed as a co-factor using two-way ANOVA, and Kruskal-Wallis used for small sample sizes in within-sex comparisons (n≤6 per group). For EPM data, the relationship between open arm entry and distance travelled was further analysed by analysis of covariance (ANCOVA). Data were analysed mainly using GraphPad Prism (v8) and IBM SPSS Statistics (v24) (two-way ANOVA). Data are presented as mean ± standard error of the mean (SEM) values. P<0.05 was considered statistically significant.

## Results

### Effect of discontinuation from 12 days of once-daily paroxetine

An exploratory experiment (6 males, 6 females per group) tested the effect of discontinuation from 12 days of once-daily paroxetine. On day 2 behaviour was assessed on the EPM. Two-way ANOVA revealed a main effect of group for time spent in the open arms (F_(2,29)_=9.050, p=0.0009) although there was also a significant sex effect (F_(1,29)_=16.85, p=0.0003) and a treatment x sex interaction (F_(2,29)_=6.357, p=0.0051). Further analysis was conducted on male and female data, separately. For male mice, there were significant main effects of group for time in the open arms (H(2)=9.789, p=0.0027; *Suppl. Fig. 1A*), open arms entries (H(2)=11.55, p=0.0005; *Suppl. Fig. 1B*) and latency for open arm entry (H(2)=6.327, p=0.0360; *Suppl. Fig. 1C*) and distance travelled (H(2)=10.96, p=0.0009; *Suppl. Fig. 1D*). Post-hoc tests showed that both discontinued and continued mice spent less time in the open arms (p=0.0309 continuation vs saline, p=0.0030 discontinuation vs saline) and made fewer open arm entries compared to saline controls (p=0.0285 continuation vs saline, p=0.0009 discontinuation vs saline). Notably, discontinued mice had an increased latency to enter the open arms (p=0.0157 vs saline) and travelled less distance (p=0.0001 vs saline). In contrast, female mice discontinued from paroxetine were not different from saline controls on any EPM parameter (*Suppl. Fig. 1A-D*), although those receiving continued paroxetine had decreased latency to open arm entry (treatment effect: H(2)=7.879, p=0.0119; *Suppl. Fig. 1C*).

On day 4 discontinued male and female mice showed no changes in spontaneous locomotion in the LMA test (*Suppl. Table 1*) and no differences on the AOF (*Suppl. Table 1*). On day 5, male discontinued mice spent less time in the light zone of the LDB (H(2)=8.667, p=0.0069; *Suppl. Fig. 1E*) with decreased distance travelled (H(2)=10.15, p=0.0021; *Suppl. Fig. 1F*) but these effects were not different from male mice on continued paroxetine (*Suppl. Fig. 1E-F*). In contrast, female mice discontinued from paroxetine actually spent more time in the light zone (H(2)=6.538, p=0.0304; *Suppl. Fig. 1E*).

Overall, results of this exploratory experiment suggest that paroxetine discontinued mice showed evidence of a behavioural effect compared to saline controls on the EPM and LDB, but this effect was not readily distingishable from continuous treatment and was apparent in males and not females.

To increase the power of this exploratory study, EPM measurements were repeated in a separate group of male mice (n=8/group additional mice) and the two male data sets were combined (n=14/group total). The pooled data confirmed that continued and discontinued mice spent less time in the open arms compared to saline controls (F_(2,38)_=6.830, p=0.0029; post-hoc p=0.0101 saline vs continuation, p=0.0011 vs saline vs discontinuation; *Fig. 1A*). Importantly, on the other EPM measures discontinued mice were significantly different to continued paroxetine as well as saline controls. Thus, discontinued mice made fewer open arms entries (F_(2,38)_=13.94, p<0.0001; post-hoc p=0.0349 vs continuation, p<0.0001 vs saline; *Fig. 1B*) and showed increased latency for open arm entry (F_(2,38)_=7.790, p=0.0015; post-hoc p=0.0201 vs continuation, p=0.0004 vs saline; *Fig. 1C*). Discontinued mice also demonstrated reduced distance travelled (F_(2,38)_=17.96, p<0.0001; post-hoc p=0.0006 vs continuation, p<0.0001 vs saline; *Fig. 1D*) as well as reduced closed arm entries (F_(2,38)_=8.546, p=0.009; *Suppl. Fig. 2*). ANCOVA revealed that changes in open arm entry were not statistically significant when co-varied with distance travelled (F_(2,37)_=1.358, p=0.270).

**Figure 1.**
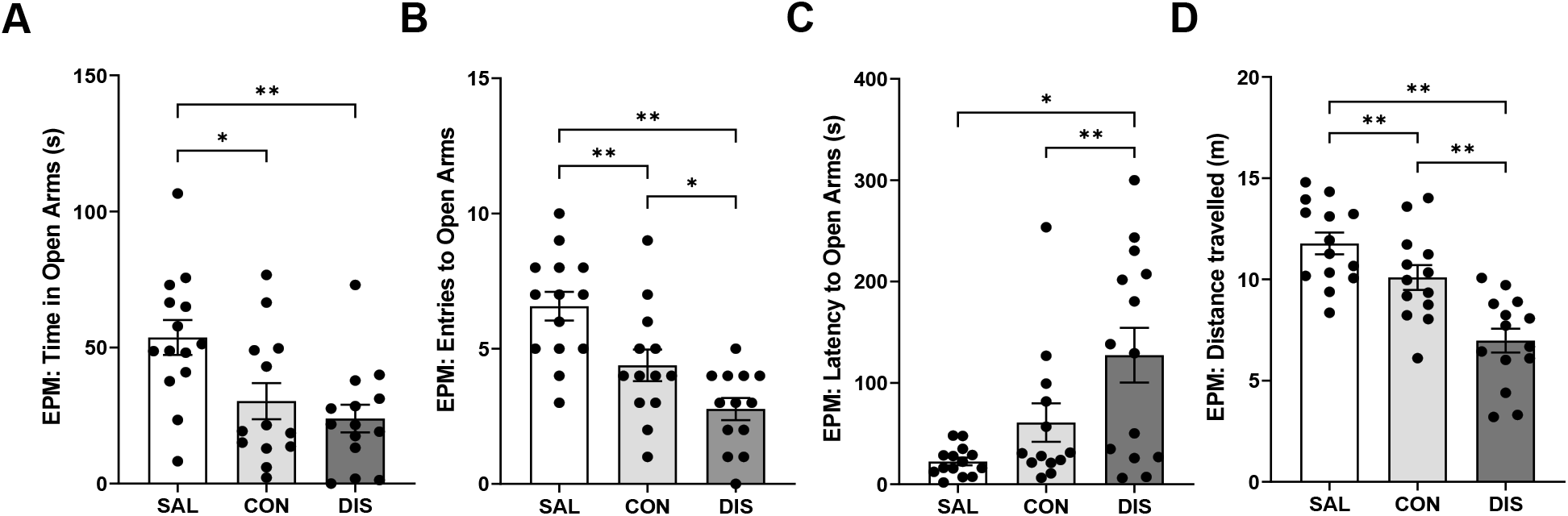
Performance on the EPM of male mice discontinued from 12 days of once-daily paroxetine. Panels show time spent in open arms (A), entries to open arms (B), latency to enter the open arms (C) and distance travelled (D). SAL, Saline (n=14); CON, Continuation (n=13, one mouse excluded due to loss of data); DIS, Discontinuation (n=14). Mean ± SEM values are shown with individual values indicated by dots. One-way ANOVA followed by post-hoc Fisher’s LSD. * p<0.05, ** p<0.01.

Notably, mice continued on paroxetine also showed decreased open arm entries (post-hoc p=0.0046 vs saline) and reduced distance travelled (post-hoc p=0.0494 vs saline) but to a lesser extent than discontinued mice (*Fig. 1*).

### Effect of discontinuation from 28 days of once-daily paroxetine

To test whether increased duration of paroxetine treatment would produce a stronger discontinuation response, male mice were treated once-daily with paroxetine for 28 days. On day 2 on the EPM, discontinued mice spent less time in the open arms compared to continued paroxetine and saline controls (F_(2,29)_=6.649, p=0.0042; post-hoc p=0.0495 vs continuation, p=0.0011 vs saline; *Fig. 2A*). Continued and discontinued mice had reduced open arm entries (F_(2,29)_=11.19, p=0.0002; post-hoc p=0.0016 continuation vs saline, p<0.0001 discontinuation vs saline; *Fig. 2B*) and increased latency for open arm entry (H(2)=13.38, p=0.0012; post-hoc p=0.0011 continuation vs saline, p=0.0021 discontinuation vs saline; *Fig. 2C*). These mice also had reduced distance travelled (F_(2,29)_=15.73, p<0.0001; post-hoc p=0.0004 continuation vs saline, p<0.0001 discontinuation vs saline; *Fig. 2D*) as well as decreased closed arm entries (F_(2,29)_=12.93, p<0.0001; *Suppl. Fig. 2*) compared to saline controls. ANCOVA revealed that changes in open arm entry were not statistically significant when co-varied with distance travelled (F_(2,28)_=1.113, p=0.343).

**Figure 2.**
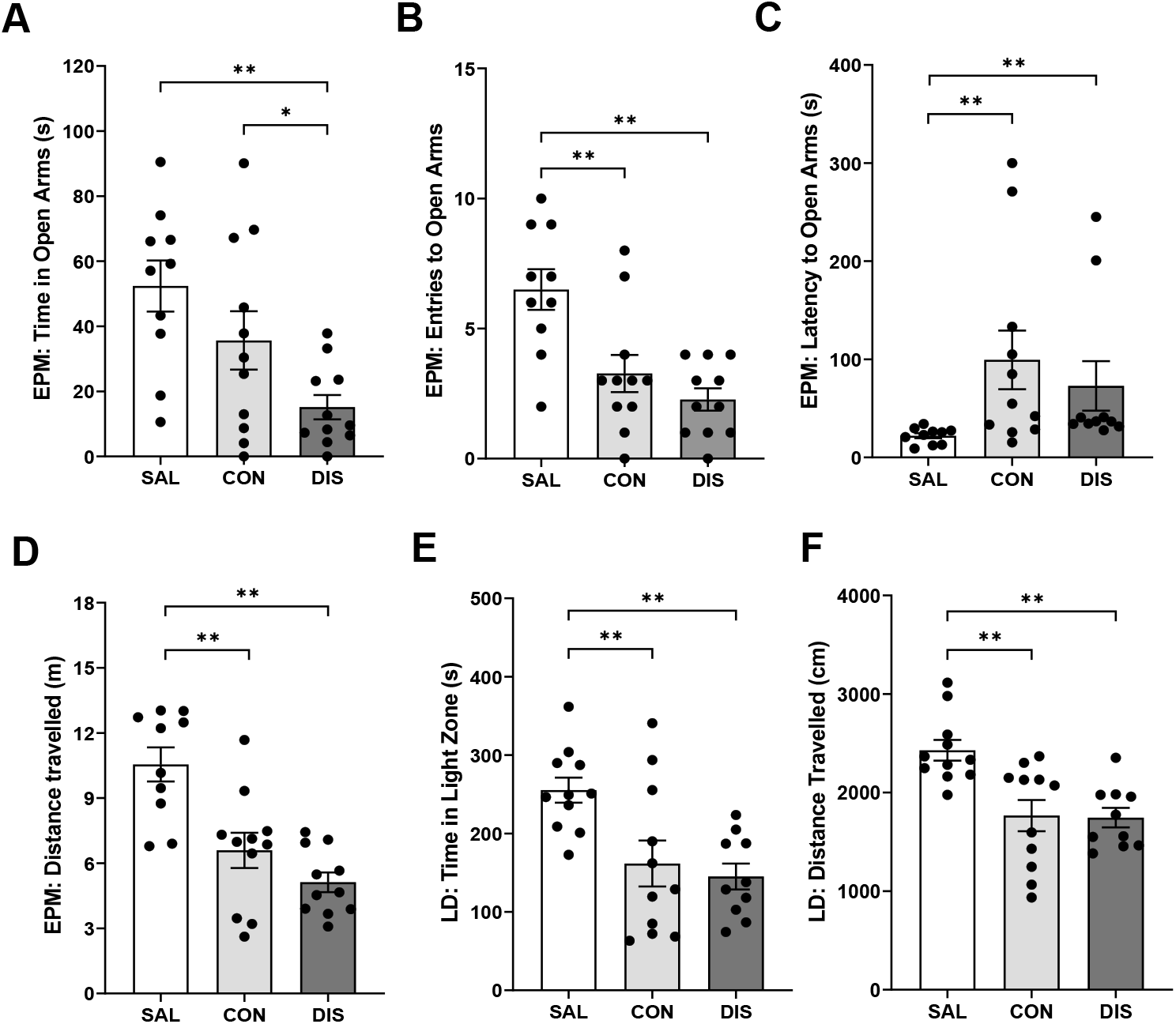
Performance on the EPM and LDB of male mice discontinued from 28 days of once-daily paroxetine. Panels show time spent in open arms (A), entries to open arms (B), latency to enter the open arms (C) and distance travelled (D) on the EPM as well as time spent in the light zone (F) and distance travelled (G) on the LDB. SAL, Saline (n=10-11); CON, Continuation (n=11); DIS, Discontinuation (n=10-11). Mean ± SEM values are shown with individual values indicated by dots. One-way ANOVA followed by post-hoc Fisher’s LSD (A-B, D-F), Kruskal-Wallis followed by post-hoc Fisher’s LSD (C). * p<0.05, ** p<0.01.

On day 4, discontinued mice had reduced activity on the LMA test compared to saline controls, as did mice on continuous treatment (F_(2,32)_=3.872, p=0.032, *Suppl. Table 2*). There was no effect of any treatment on performance on the AOF compared to saline controls (*Suppl. Table 2*). On day 5, discontinued mice spent less time in the light zone of the LDB (F_(2,29)_=7.468, p=0.0024; post-hoc p=0.0014 vs saline; *Fig. 2E*) and had decreased distanced travelled (F_(2,29)_=9.782, p=0.0006; post-hoc p=0.0006 vs saline; *Fig. 2F*) compared to saline controls, but these effects were not different from mice on continued paroxetine (time in light zone: post-hoc p=0.0045 continuation vs saline; distance travelled: post-hoc p=0.0007 continuation vs saline; *Fig. 2*).

Overall, these results add evidence that 28 day paroxetine generates a discontinuation effect on the EPM although this was not obviously a greater and longer lasting than 12 day paroxetine treatment.

### Effect of discontinuation from 7 days of once-daily paroxetine

To investigate the duration of treatment with paroxetine required to produce discontinuation effects, male mice received once-daily paroxetine injections for 7 days. On day 2 following treatment cessation, discontinued mice did not perform differently on the EPM compared to saline controls or mice on continuous paroxetine in terms of time spent in the open arms (F_(2,33)_=0.1256, p=0.8824, *Fig. 3A*), open arm entries (F_(2,33)_=0.6936, p=0.5069; *Fig. 3B*), latency to open arm entry (H(2)=1.610, p=0.4471; *Fig. 3C*), distance travelled (F_(2,33)_=1.262, p=0.2964; *Fig. 3D*) or closed arm entries (F_(2,33)_=1.734, p=0.1923; *Suppl. Fig. 2*).

**Figure 3.**
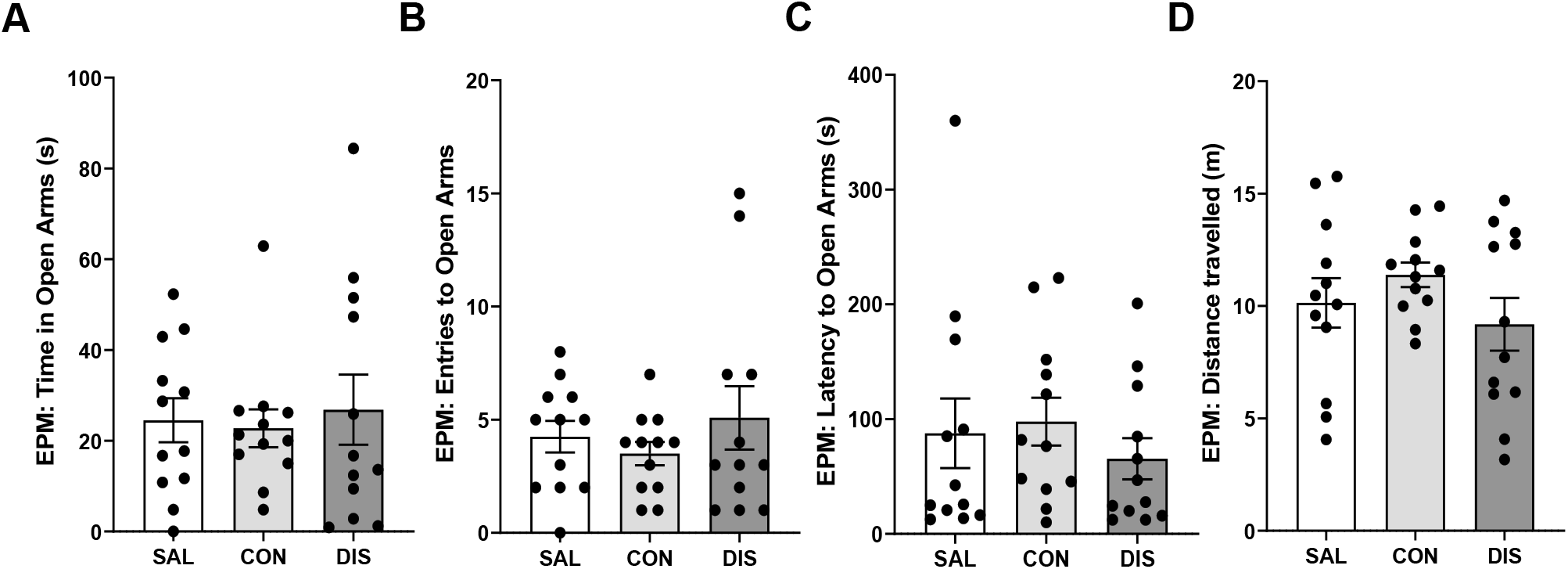
Performance on the EPM of male mice discontinued from 7 days of once-daily paroxetine. Panels show time spent in open arms (A), entries to open arms (B), latency to enter the open arms (C) and distance travelled (D). SAL, Saline (males n=12); CON, Continuation (males n=12); DIS, Discontinuation (males n=12). Mean ± SEM values are shown with individual values indicated by dots. One-way ANOVA followed by post-hoc Fisher’s LSD (A-B, D), Kruskal-Wallis followed by post-hoc Fisher’s LSD (C).

These results show that paroxetine treatment for 7 days was not sufficient to induce a discontinuation response on the EPM.

### Effect of discontinuation from 12 days of twice-daily paroxetine

To test whether increasing the frequency of paroxetine treatment (and hence increasing daily dose) would produce a more pronounced discontinuation effect on the EPM, mice were treated twice-daily with paroxetine for 12 days. This experiment also tested the possibility that mice continuously treated with paroxetine may have been experiencing discontinuation effects, in that up to this point all ‘continuous’ treatment groups were tested 18 h after the last injection. Here mice were tested 6 h after the last injection so that plasma levels of paroxetine likely remained high. This experiment also examined the possibility that discontinuation from paroxetine would affect fear-learning, which is reported to be sensitive to manipulations of the 5-HT transporter (e.g. (Lima et al., 2019)).

On day 2 following treatment cessation, discontinued mice showed reduced time on the open arms (F_(2,33)_=10.83, p=0.0002; post-hoc p<0.0001 vs saline; *Fig. 4A*) and reduced open arm entries (F_(2,33)_=3.482, p=0.0424; post-hoc p=0.0141 vs saline; *Fig. 4B*), albeit with unchanged latency for open arm entry (F_(2,33)_=2.174, p=0.3373; *Fig 4C*). Discontinued mice also demonstrated reduced distance travelled (F_(2,33)_=7.469, p=0.0021; post-hoc p=0.0086 vs continuation, p=0.0008 vs saline; *Fig. 4D*) and decreased closed arm entries (F_(2,33)_=8.466, p=0.0011; *Suppl. Fig. 2*). Again, ANCOVA revealed that changes in open arm entry were not statistically significant when co-varied with distance travelled (F_(2,28)_=0.123, p=0.884).

**Figure 4.**
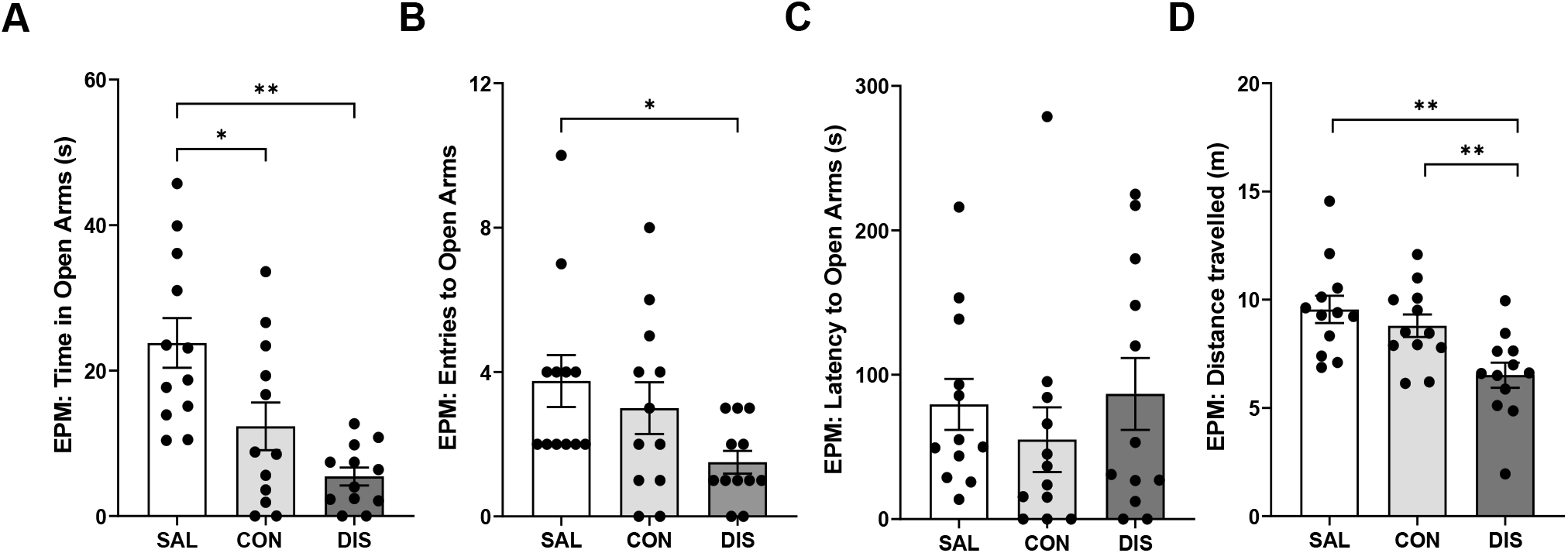
Performance on the EPM male mice discontinued from 12 days of twice-daily paroxetine. Panels represent time spent in open arms (A), entries to open arms (B), latency to enter the open arms (C) and distance travelled (D). SAL, Saline (n=12); CON, Continuation (n=12); DIS, Discontinuation (n=12). Mean ± SEM values are shown with individual values indicated by dots. One-way ANOVA followed by post-hoc Fisher’s LSD (A-B, D), Kruskal-Wallis followed by post-hoc Fisher’s LSD (C). * p<0.05, ** p<0.01.

In comparison, mice receiving continued paroxetine also spent less time in open arms (post-hoc p=0.0070 vs saline; *Fig. 4A*) even when tested 6 h after the last injection, suggesting these changes are not in themselves discontinuation effects.

Importantly, given the above evidence that discontinued mice had reduced motor activity on the EPM, on day 3 mice were tested for locomotor changes in a separate LMA test. Neither discontinued mice nor mice receiving continued paroxetine were different from saline controls in on the LMA test (F_(2,33)_=0.073, p=0.929). There was also no effect of any treatment on activity on the AOF (*Suppl. Table 3*).

On days 4 and 5 following cessation of paroxetine treatment, mice were assessed for fear conditioning (training on day 4, testing on day 5). All mice exhibited increased freezing with the onset of the conditioned stimulus. However, neither discontinued mice nor mice on continued treatment were different from saline controls in terms of freezing to tone on the training day or in their response to the conditioned tone during recall on the test day (*Suppl. Table 3*).

These results add further evidence that paroxetine-treated mice produces a discontinuation response on the EPM, but that increased treatment frequency (and thereby dose) did not lead to a more clear-cut and longer-lasting effects. Furthermore, separate LMA testing found no evidence of a general reduction in motor activity in the discontinued mice.

### Effect of discontinuation from 12 days of twice-daily citalopram

A final experiment determined the effects of discontinuation from 12 days of twice-daily treatment with another SSRI, citalopram. On day 2 following treatment cessation, discontinued mice showed increased latency to enter the open arms (H(2)=7.220, p=0.0221; post-hoc p=0.0078 vs saline; *Fig. 5C*) and a trend effect for reduced number of open arm entries (F_(2,51)_=1.751, p=0.1839; post-hoc p=0.0862 vs continuation; *Fig. 5B*) although time spent in the open arms was not different between groups (F_(2,51)_=0.5349; 0.5890; *Fig. 5A*). Discontinuation from citalopram had no effect on distance travelled (F_(2,51)_=1.949, p=0.1529; *Fig. 5D*) or closed arm entries (F_(2,27)_=1.323, p=0.2830; *Suppl. Fig. 2*). Unlike mice treated continuously with paroxetine, mice on continuous citalopram (tested 6 h after the last injection) were not different from saline controls on the EPM (*Fig. 5*).

**Figure 5.**
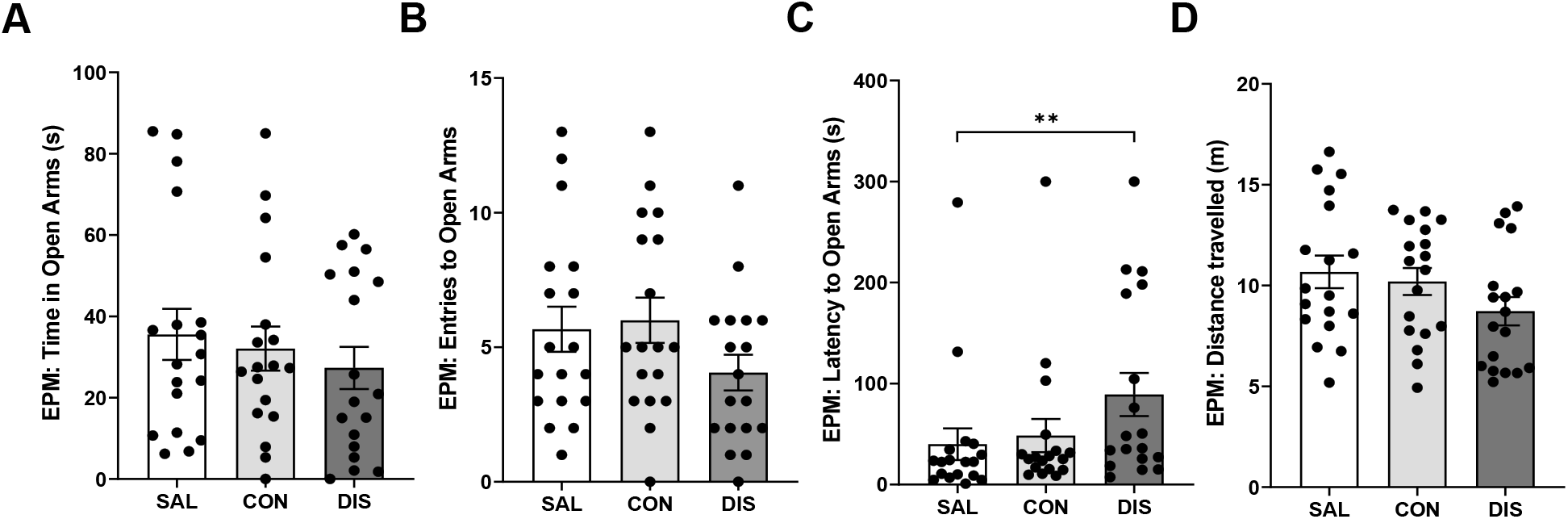
Performance on the EPM of male mice discontinued from 12 days of twice-daily citalopram. Panels represent time spent in open arms (A), entries to open arms (B), latency to enter the open arms (C) and distance travelled (D). SAL, Saline (n=18); CON, Continuation (n=18); DIS, Discontinuation (n=18). Mean ± SEM values are shown with individual values indicated by dots. One-way ANOVA followed by post-hoc Fisher’s LSD (A-B, D), Kruskal-Wallis followed by post-hoc Fisher’s LSD (C). * p<0.05, ** p<0.01.

Mice discontinued from citalopram were also not different from either saline controls or mice maintained on citalopram on the LMA and AOF tests (day 3 of discontinuation), nor on any parameter measured during the training (day 4) or recall (day 5) stages of the fear conditioning paradigm (*Suppl. Table 4*).

These results suggest that as with paroxetine, citalopram produced a discontinuation response on the EPM albeit a modest one.

## Discussion

Abrupt cessation of a course of treatment with an SSRI in humans is often associated with a discontinuation syndrome comprising unpleasant centrally and peripherally-mediated changes, but the effects of SSRI discontinuation in animals is little studied. The present study determined the effect of SSRI discontinuation in mice with a focus both on anxiety-like behaviour, which is a common discontinuation symptom, and paroxetine which is particularly problematic in patients. A main finding was that 2 days following discontinuation from repeated administration of paroxetine mice showed differences in behaviour on the EPM compared to mice maintained on drug, as well as saline controls. A modest discontinuation effect on the EPM was also detected with citalopram. Attempts to optimise treatment protocols found that paroxetine exposure for more than a week was required to elicit a discontinuation effect, and this was not obviously strengthened with increasing duration or dose of treatment. Data from other anxiety tests showed evidence of persistent behavioural effects in mice up to 5 days after paroxetine discontinuation; however, these effects were relative to saline controls and not mice maintained on drug (specifically paroxetine). Finally, the discontinuation effect of paroxetine was sexually-dymorphic in that, at least under the present conditions, the response was detected in male and not female mice. Overall, the current data provide the first evidence for the emergence of a discontinuation effect in male mice within 2 days of cessation of SSRI treatment although the experiments were unable to resolve the duration of this effect.

The EPM is widely used to assess anxiety levels in rodents, and has been previously used to detect increased anxiety evoked by discontinuation of other drugs including psychostimulants and alcohol, as well as for detecting changes in 5-HT function (see Introduction). The majority of these studies prioritise two EPM indices of anxiety: open arm entries and time spent on the open arms, which are closely correlated parameters (Lister, 1990). Here, one or other of these parameters were found to be decreased in SSRI discontinued mice compared to those maintained on treatment. Detection of anxiety-like behaviours on the EPM can be confounded by non-specific changes in motor activity (Lister, 1990; Handley and McBlane, 1993; Weiss et al., 1998), and in the present study SSRI discontinuation was often associated with reduced distance travelled on the EPM which may have impacted on open arm exploration. ANCOVA can be used to help determine the influence of changes in distance travelled on open arm exploration parameters, but here these changes could not be separated by this statistical approach. However, the impact of ANCOVA is limited as pointed out elsewhere (Lister, 1990). Thus, in the context of an anxiogenic environment, a decrease in distance travelled would be consistent with an increase in anxiety since behavioural inhibition is a key component of the anxiety response (Gray, 1982; McNaughton and Gray, 2000). Thus, these EPM parameters are very likely to co-vary.

To further explore the possibility that reduced open arm exploration was simply due to decrease in general motor activity, we carried out separate locomotor assessments in a low anxiety, home cage-like environment as previously recommended (Lister, 1990). This separate assessment found little evidence that mice undergoing SSRI discontinuation exhibited a reduction in general motor activity (with the exception of the 28 day paroxetine, see below). Thus, decreased locomotion on the EPM appeared to be context-specific and plausibly arose from increased anxiety levels due to exposure to a novel, anxiogenic environment rather than decreased locomotor activity *per se*. Nevertheless, without further investigation we cannot exclude the possibility that the elevated anxiety measures in discontinued mice is not genuine anxiety but instead linked to other unpleasant experiences of discontinuation including somatic changes that feature in the clinical syndrome, which in itself would be of great interest. Indeed, discontinuation from 28 day paroxetine were confounded by reduced locomotion in separate assessments suggesting the emergence of a syndrome which may be driven by hypolocomotion.

The evidence of altered EPM performance within 2 days of discontinuation from paroxetine and citalopram is consistent with the short half-lives of these drugs in rodents (6.3 h and 1.5 h, respectively) (Kreilgaard et al., 2008; Fredricson Overø, 1982), and an abrupt fall in brain levels of the drug once drug administration has stopped. Moreover, a rapid appearance of a discontinuation effect on the EPM is consistent with earlier evidence of heightened startle responsivity in rats 2 days after discontinuation from repeated treatment with citalopram (~24 mg/kg per day for 15 days) (Bosker et al., 2010). Our finding also fits with the clinical picture that SSRIs like paroxetine, which have a short half-life in humans, are particularly problematic with patients often experiencing discontinuation effects within two days of treatment cessation (Rosenbaum et al., 1998; Michelson et al., 2000; Fava et al., 2015). Although plasma levels of paroxetine and citalopram were not measured in the present experiments, previous studies report that the administration of paroxetine or citalopram to rodents at the doses used here produces plasma drugs levels broadly in the therapeutic range, and that fall quickly once administration ceases. For example, rat plasma levels of citalopram (121 ng/ml) were reported to fall around 8-fold within 24 h of stopping administration (24 mg/kg per day for 15 days) and were at the limits of detection at 48 h (Bosker et al., 2010). Similarly, rat plasma levels of paroxetine (411 ng/ml) fell by a similar degree 48 h after cessation of paroxetine (10 mg/kg for 21 days) and were undetectable after 96 h of washout (Benmansour et al., 1999).

A clinical feature of SSRI discontinuation is that in some patients the effect can endure for several weeks or even months (Fava et al., 2007; Davies and Read, 2019). The present study applied a battery of different anxiety tests to determine whether effects of SSRI discontinuation persist since animals show an adaptive response following repeated testing on the EPM test and other anxiety tests (either due to habituation or behavioural changes from anxiety to acquired fear (File, 2001)). This test battery also determined the generality of our findings across assays with different sensorimotor and motivational demands. However, results were somewhat inconclusive. For example, mice discontinued from paroxetine (eg. once-daily for 12 days) showed evidence of increased anxiety on the LDB on day 5 compared to saline controls, but it was not statistically different from mice receiving continuous paroxetine in this test. These data potentially accord with a short-lasting anxiogenic effect of SSRI discontinuation in mice that may dissipate after 48 h. However, a previous study found an enhanced acoustic startle response in rats discontinued from citalopram compared to those still receiving treatment that persisted for at least 7 days after the last dose (Bosker et al., 2010). The current data do not preclude the persistence of discontinuation effects as it is possible that the use of a battery of tests over consecutive days obscured the detection of genuine discontinuation versus continuation effects due to prior test exposure.

Here, a discontinuation response to paroxetine was detected after 12 and 28 days but not after 7 days of continuous drug treatment, suggesting the involvement of a neuroadaptive mechanism. This timing accords with findings in clinical studies that taking an SSRI for a week is not sufficient to produce discontinuation symptoms (Yonkers et al., 2015). Further, randomised control trials show that discontinuation symptoms already emerge after a few weeks of SSRI treatment (Baldwin et al., 2006). The nature of the neuroadaptive responses underpinning SSRI discontinuation is unknown. Although much is established regarding how the 5-HT systems adapts during a course of SSRI administration (Sharp, 2013; Artigas, 2013), remarkably few studies have investigated how 5-HT transmission changes when chronic SSRI administration is immediately stopped. Two earlier studies reported decreased 5-HT synthesis and metabolism in post-mortem rat brain tissue for up to 3 days following withdrawal from citalopram and fluoxetine (Trouvin et al., 1993; Bosker et al., 2010). On the other hand, SSRI discontinuation likely leaves 5-HT neurons in a state of reduced 5-HT_1A_ autoreceptor sensitivity (desensitisation), which theoretically would push the 5-HT system in the opposite direction and increase 5-HT neuron firing and 5-HT release (Richardson-Jones et al., 2010). Indeed, Trouvin and colleagues (1993) reported an overshoot in brain 5-HT metabolite levels 7-21 days following discontinuation from fluoxetine. Although the delayed timing of this 5-HT overshoot is out of keeping with the current findings, it is in line with the other lines of research reporting that increased anxiety is associated with increased 5-HT (Briley et al., 1990; Handley and McBlane, 1993; Ohmura et al., 2020).

Neuroadaptive changes in certain postsynaptic 5-HT receptors as well as the 5-HT transporter itself are also reported following genetic knockout or pharmacological inhibition of the 5-HT transporter (Jennings et al., 2012; Benmansour et al., 1999) and might rebound during discontinuation. Thus, contrary to earlier speculations of either a hyper- or hyposerotonergic state underpinning the symptoms of SSRI discontinuation syndrome (Blier and Tremblay, 2006; Renoir, 2013), a potentially more complex and dynamic picture emerges of decreased synaptic availability of 5-HT, elevated 5-HT cell firing, altered postsynaptic 5-HT receptors, and overall destabilised (and dysfunctional) 5-HT signalling and related 5-HT-induced changes in neural plasticity. Although as a <u>drug class</u> SSRIs are pharmacologically selective, 5-HT has multiple interactions with other transmitter systems, including the co-release of glutamate from 5-HT neurons and the 5-HT regulation of noradrenergic, dopaminergic and GABAergic circuits (Sharp and Barnes, 2020). Thus, it is plausible that such transmitters also contribute to the effects of SSRI discontinuation. The key role of noradrenaline in the effects of opiate withdrawal (eg. Kosten and Baxter, 2019) illustrates the potential link between discontinuation effects and other neurotransmitter systems. Finally, numerous studies have reported enhanced neuroplasticity and structural changes following repeated SSRI treatment which are thought to be responsible for their therapeutic effects (Castrén and Hen, 2013; Harmer et al., 2017). For example, paroxetine induced an adaptive increase in hippocampal neurogenesis following several weeks of treatment in mice (Elizalde et al., 2010), which plausibly relates to the behavioural effects of paroxetine discontinuation observed here. However, it is unlikely that such changes reverse immediately following SSRI discontinuation. Nonethless, the lack of information on the timing of changes in plasticity such as neurogenesis upon SSRI discontinuation means this possibility cannot currently be excluded.

Our initial exploratory study did not detect an effect of discontinuation from paroxetine (once-daily, 12 days) in female mice. Further investigation of this finding is required as there are many sexual dymorphisms reported in mice, some of which offer a plausible explanation for the male-female difference found here. For example, female mice metabolise fluoxetine more effectively than males through increased activity of cytochrome P_450_ enzymes (Hodes et al., 2010). Since paroxetine is also metabolised by these enzymes, in the present study the paroxetine dosing regimen may not have been sufficiently high for female mice. Also, evidence of sexual dymorphism in the 5-HT system is reported for mice (Renoir et al., 2011), in particular female mice were found to exhibit reduced 5-HT transporter mRNA compared to males. If this difference translated to the female mice used here, it would account for their reduced sensitivity to paroxetine. The stage of the estrous cycle of mice was also found to influence the behavioural response to fluoxetine treatment on tests of anxiety, with mice in the diestrus phase of the cycle being less sensitive the drug (Yohn et al., 2020). Furthermore, we cannot exclude the possibility that these differences reflect different homecage experiences of male and female mice, particularly in terms of aggressive behaviours and social dominance hierarchies. Nonetheless, there is evidence from clinical studies that men are more likely to report discontinuation symptoms following SSRI cessation (Coupland et al., 1996), so the current findings may reflect a the greater propensity of males to experience SSRI discontinuatuion syndrome rather than pitfalls of the preclinical model.

Finally, it is interesting to note that in the present experiments, mice *maintained* on paroxetine showed evidence of increased anxiety on the EPM and LDB. One possible explanation is that discontinuation effects are occurring even in these animals because in some experiments mice were tested 18 h after the last injection of paroxetine (see *Table 1*) when drug levels are likely to have fallen. However, since mice tested 6 h after the last injection still showed evidence of increased anxiety, discontinuation is not the likely cause. Another explanation is that increased anxiety is due to continuing blockade of the 5-HT transporter, and the consequent increase in synaptic 5-HT. In support of this idea, previous studies with mice reported that continuous treatment with certain SSRIs is anxiogenic on the EPM (Oh et al., 2009; Venzala et al., 2012; Turcotte-Cardin et al., 2019). Ourselves and others have also reported that 5-HT transporter knockout mice show heightened anxiety on the EPM and other anxiety tests (Ansorge et al., 2004; Line et al., 2011). However, other findings argue against this position. Firstly, the high anxiety phenotype of the 5-HT transporter knockout mice may in part involve a neurodevelopmental mechanism (Ansorge et al., 2004). Secondly, previous studies on the effect of repeated administration of paroxetine to mice on EPM performance do not consistently detect increased anxiety (Goeldner et al., 2005; Elizalde et al., 2008; Thoeringer et al., 2010; Guilloux et al., 2011), but mouse strain difference are one potential confound (Jin et al., 2017). Thirdly, the current study found that mice continuously treated with citalopram (twice-daily for 12 days) did not show an anxiogenic response on the EPM. Thus, on balance, available data suggest that the anxiogenic effect of continuous paroxetine detected here is not easily explained by either discontinuation itself or a simple 5-HT-related mechanism. Alternative explanations include a role for other specific pharmacological effects of paroxetine (muscarinic receptor antagonism, noradrenaline reuptake blockade).

In conclusion, the present study reports evidence of a discontinuation response in mice within 48 h of cessation from a course of SSRI treatment, that likely involves a neuroadaptive mechanism. This discontinuation response had features consistent with anxiety-provoked behavioural inhibition rather than a general reduction in motor activity, which correlates with findings of increased anxiety in patients within days of stopping a course of SSRI therapy. SSRI discontinuation in mice may provide a useful model to aid the investigation of the neurobiological mechanisms involved.

## Acknowledgements

The authors disclosed receipt of the following financial support for the research, authorship, and/or publication of this article: HMC was funded by a Wellcome Trust Four-Year PhD Studentship in Basic Science (219982/Z/19/Z). For the purpose of open access, the author has applied a CC BY-ND public copyright licence to any Author Accepted Manuscript version arising from this submission.

