## Supplementary Figure1 for "Effect of SSRI discontinuation on anxiety-like behaviours in mice"

### Supplementary information

#### Supplementary Figure 1

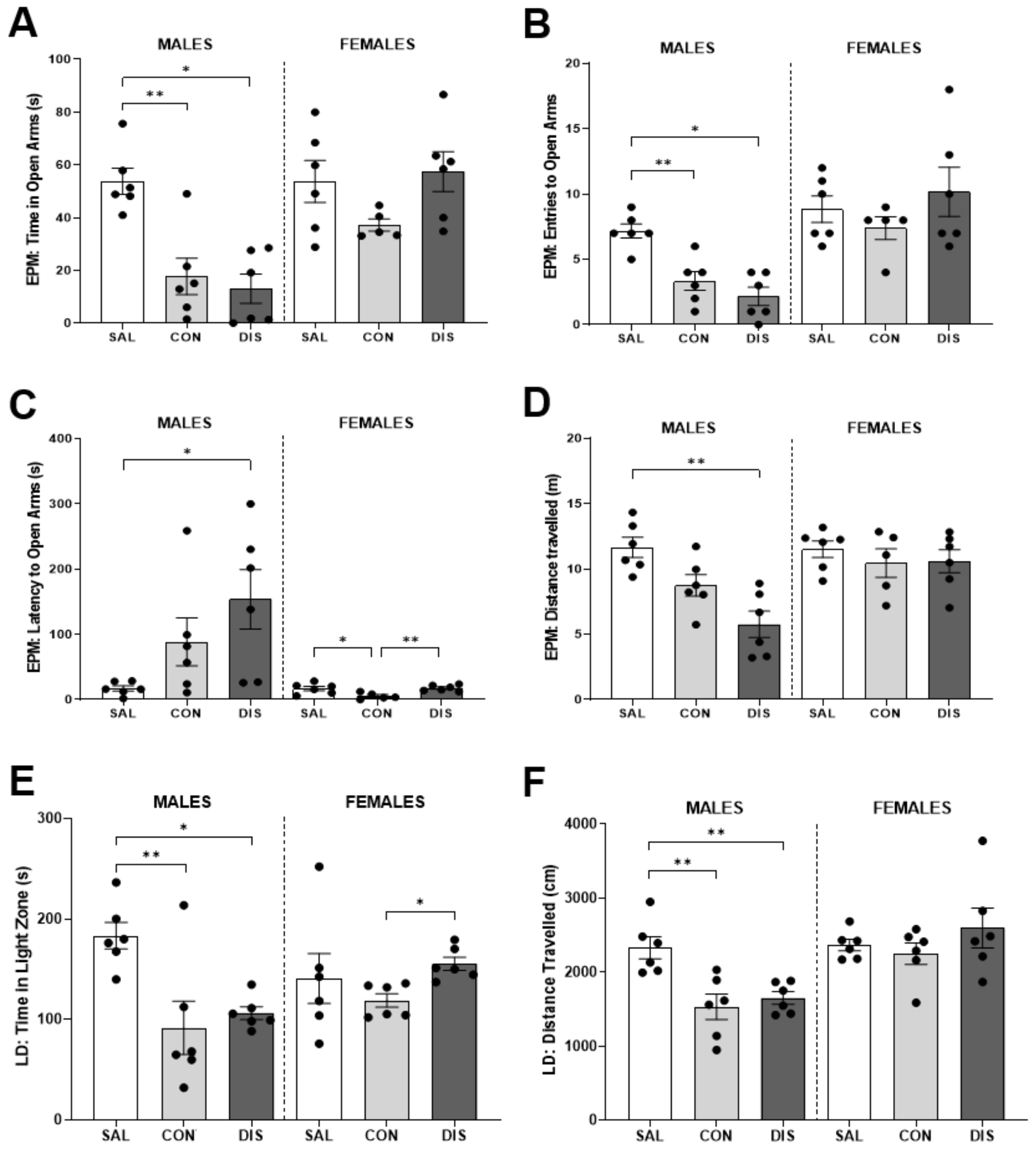


Effect of discontinuation from 12 days of once-daily paroxetine treatment in male and female mice on the elevated plus maze (EPM; 300 s) and the light/dark box (LDB; 300 s). Bars represent the mean ± SEM values for time spent in open arms (males: p=0.0309 SAL vs CON, p=0.0030 SAL vs DIS) (A), entries to open arms (males: p=0.0285 SAL vs CON, p=0.0009 SAL vs DIS) (B), latency to enter the open arms (males: p=0.0157 SAL vs DIS; females: p=0.0355 SAL vs CON, p=0.0189 CON vs DIS) (C) and distance travelled (males: p=0.0001 SAL vs DIS) (D) on the EPM, and time spent in the light zone (males: p=0.0058 SAL vs CON, p=0.0231 SAL vs DIS; females: p=0.00110 CON vs DIS) (F) and distance travelled (males: p=0.0069 SAL vs CON, p=0.0049 SAL vs

DIS) (G) on the LDB. SAL, Saline (males n=6, females n=6); CON, Continuation (males n=6, females n=5, one female mouse excluded due to technical issues during testing); DIS, Discontinuation (males n=6, females n=6). Individual values are indicated by dots. Kruskal-Wallis followed by post-hoc Fisher's LSD, * $p<0.05$, ** $p<0.01$.

**Supplementary Figure 2**

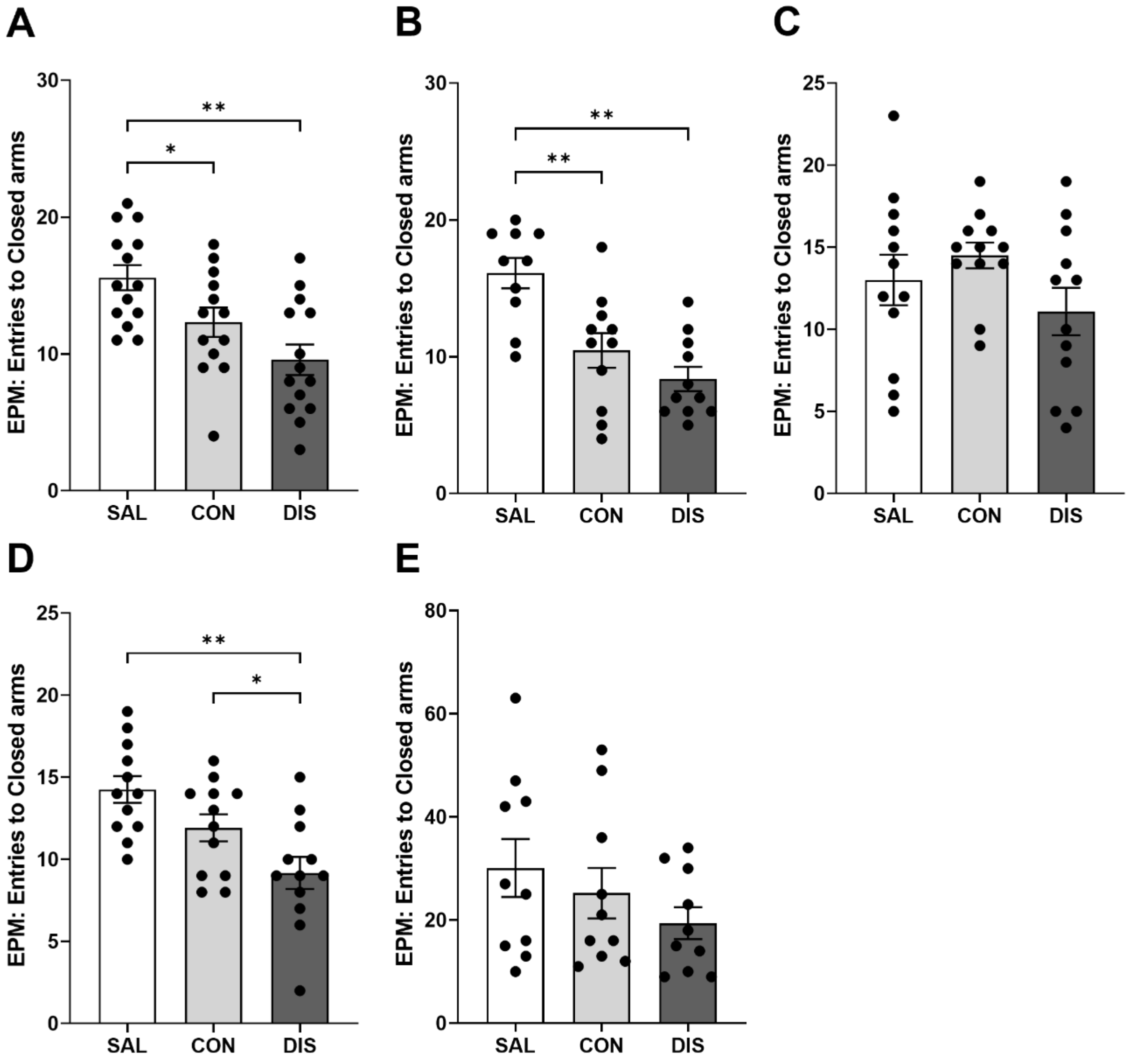


Effect of discontinuation from paroxetine or citalopram treatment in male mice on the closes arm entries on the elevated plus maze (EPM; 300 s). Bars represent the mean ± SEM values for number of entries to the closed arms following discontinuation from 12-days once-daily paroxetine (p=0.0336 SAL vs CON, p=0.0002 SAL vs DIS) (A), 28-days of once-daily paroxetine (p=0.0012 SAL vs CON, p<0.0001 SAL vs DIS) (B), 7-days once-daily paroxetine (C), 12-days twice-daily paroxetine (p=0.0002 SAL vs DIS, p=0.0331 CON vs DIS) (D) and 12-days twice-daily citalopram (E). SAL, Saline (n=10-14); CON, Continuation (n=10-14); DIS, Discontinuation (n=10-14). Individual values are indicated by dots. One-way ANOVA followed by post-hoc Fisher's LSD, * p<0.05, ** p<0.01.

**Supplementary Table 1**

Effect of Saline (SAL), Continuation (CON) or Discontinuation (DIS) treatments with 12-days once-daily paroxetine in male and female mice. LMA, locomotor activity (60 min); AOF, aversive open field (600 s). Each value is a mean ± S.E.M.

| | MALES (n=6/group) | | | | FEMALES (n=6/group) | | | |
|---|---|---|---|---|---|---|---|---|
| | Kruskal | Group (mean ± SEM) | | | Kruskal | Group (mean ± SEM) | | |
| **Parameter** | **Wallis** | **SAL** | **CON** | **DIS** | **Wallis** | **SAL** | **CON** | **DIS** |
| *LMA* | | | | | | | | |
| Total beam breaks | H(2)=1.906 p=0.408 | 2325 ± 239 | 1974 ± 163 | 1857 ± 147 | H(2)=4.433 p=0.108 | 2677 ± 140 | 2314 ± 181 | 3274 ± 461 |
| Total rearings | H(2)=2.854 p=0.251 | 175 ± 17 | 165 ± 10 | 191 ± 5 | H(2)=1.556 p=0.484 | 162 ± 19 | 179 ± 14 | 196 ± 13 |
| *AOF* | | | | | | | | |
| Time in centre (s) | H(2)=0.737 p=0.715 | 13.7 ± 2.5 | 24.2 ± 9.3 | 11.2 ± 3.0 | H(2)=0.924 p=0.652 | 11.1 ± 1.5 | 14.9 ± 4.6 | 22.3 ± 7.5 |
| Distance travelled (m) | H(2)=3.591 p=0.170 | 50.1 ± 4.5 | 40.3 ± 5.1 | 45.5 ± 3.7 | H(2)=0.947 p=0.640 | 58.8 ± 3.7 | 54.0 ± 6.6 | 74.5 ± 16.2 |

**Supplementary Table 2**

Effect of Saline (SAL), Continuation (CON) or Discontinuation (DIS) treatments with 28-day once-daily paroxetine in male mice. LMA, locomotor activity (60 min); AOF, aversive open field (600 s); LDB, light/dark box (600 s). 1 mouse excluded from the LMA onwards due to receiving the wrong injection the night before; 1 mouse excluded from the LMA due to issues with the photobeam box during testing. Each value is a mean ± S.E.M., and the statistical outcomes of pairwise comparisons were: SAL vs DIS *p<0.05; SAL vs CON †p<0.05, ††p<0.01.

| | | **Paroxetine (n=12/group)** | | |
|---|---|---|---|---|
| | **ANOVA/** | | **Group (mean ± SEM)** | |
| | **Kruskal Wallis** | **SAL** | **CON** | **DIS** |
| *LMA* | | | | |
| Total beam breaks | $F_{(2,31)}$=3.872 p=0.032 | 2522 ± 154 | 2061 ± 148 † | 2044 ± 110 * |
| Total rearings | $F_{(2,31)}$=0.336 p=0.717 | 167.2 ± 14.5 | 181.5 ± 8.7 | 169.3 ± 15.5 |
| *AOF* | | | | |
| Time in centre (s) | $F_{(2,32)}$=2.327 p=0.114 | 11.9 ± 1.0 | 8.5 ± 1.6 | 7.8 ± 1.7 |
| Distance travelled (m) | $F_{(2,32)}$=10.01 p<0.001 | 64.6 ± 4.7 | 40.1 ± 3.0 †† | 49.8 ± 4.0 * |

**Supplementary Table 3**

Effect of Saline (SAL), Continuation (CON) or Discontinuation (DIS) treatments with 12-days twice-daily paroxetine in male mice. LMA, locomotor activity (60 min); AOF, aversive open field (600 s); FC, fear conditioning; CS, conditioned stimulus. 1 SAL and 1 CON mouse excluded due to technological issues on training day. Each value is a mean ± S.E.M., and the statistical outcomes of pairwise comparisons were: SAL vs DIS *p<0.01; SAL vs CON †p<0.05.

| | | **Paroxetine (n=11-12/group)** | | |
|---|---|---|---|---|
| | **ANOVA/ Kruskal Wallis** | **Group (mean ± SEM)** | | |
| | | **SAL** | **CON** | **DIS** |
| *LMA* | | | | |
| Total beam breaks | $F_{(2,33)}$=0.073, p=0.929 | 1913.7 ± 161.7 | 1906.8 ± 137.4 | 1839.5 ± 134.3 |
| Total rearings | $F_{(2,33)}$=1.293, p=0.288 | 190.4 ± 10.4 | 170 ± 10.4 | 173.1 ± 6.3 |
| *AOF* | | | | |
| Time in centre (s) | $F_{(2,32)}$=2.167, p=0.1306 | 6.6 ± 0.9 | 3.8 ± 1.0 | 5.7 ± 1.0 |
| Distance travelled (m) | $F_{(2,33)}$=5.560, p=0.0083 | 28.3 ± 1.8 | 21.6 ± 1.7 † | 21.4 ± 1.5 * |
| *FC* | | | | |
| Training day | | | | |
| Δ Freezing (%) | $F_{(2,31)}$=2.549, p=0.0944 | 5.6 ± 1.8 | -0.6 ± 1.8 | 2.4 ± 2.1 |
| Test day | | | | |
| Pre-CS freezing (%) | $F_{(2,31)}$=0.2451, p=0.7842 | 19.2 ± 6.6 | 25.6 ± 8.1 | 22.4 ± 4.0 |
| Post-CS freezing (%) | $F_{(2,31)}$=0.1248, p=0.8831 | 39.3 ± 7.5 | 38.2 ± 8.9 | 43.1 ± 5.6 |
| Δ Freezing (%) | $F_{(2.31)}$=0.3106, p=0.7353 | 19.9 ± 5.9 | 14.6 ± 4.5 | 19.03 ± 4.6 |

**Supplementary Table 4**

Effect of Saline (SAL), Continuation (CON) or Discontinuation (DIS) treatments with 12-days twice-daily citalopram in male mice. LMA, locomotor activity (60 min); AOF, aversive open field (600 s); FC, fear conditioning. Each value is a mean ± S.E.M.

| | | **Citalopram (n=10/group)** | | |
|---|---|---|---|---|
| | **ANOVA/ Kruskal Wallis** | **Group (mean ± SEM)** | | |
| | | **SAL** | **CON** | **DIS** |
| *LMA* | | | | |
| Total beam breaks | $F_{(2,27)}$=0.7999, p=0.4598 | 2491.0 ± 131.1 | 2190.0 ± 261.5 | 2298.0 ± 260.4 |
| Total rearings | $F_{(2,27)}$=1.6710, p=0.0276 | 202.3 ± 6.6 | 156.1 ± 16.2 | 171.4 ± 10.0 |
| *AOF* | | | | |
| Time in centre (s) | $F_{(2,27)}$=0.4738, p=0.6279 | 9.1 ± 1.2 | 9.3 ± 1.0 | 8.9 ± 1.0 |
| Distance travelled (m) | $F_{(2,27)}$=0.6227, p=0.5440 | 37.4 ± 3.0 | 32.3 ± 2.3 | 32.3 ± 3.1 |
| *FC* | | | | |
| Training day | | | | |
| Δ Freezing (%) | $F_{(2,27)}$=0.9676, p=0.3928 | 6.8 ± 2.4 | 12.8 ± 3.1 | 8.0 ± 4.0 |
| Test day | | | | |
| Pre-CS freezing (%) | $F_{(2,27)}$=2.247, p=0.1073 | 17.0 ± 5.2 | 43.5 ± 9.0 | 26.3 ± 9.9 |
| Post-CS freezing (%) | $F_{(2,27)}$=2.932, p=0.0704 | 60.5 ± 6.0 | 75.8 ± 5.3 | 53.3 ± 26.6 |
| Δ Freezing (%) | $F_{(2,27)}$=1.215, p=0.3124 | 43.5 ± 6.5 | 33.3 ± 7.1 | 27.0 ± 8.9 |
